# High-Precision Automated Reconstruction of Neurons with Flood-filling Networks

**DOI:** 10.1101/200675

**Authors:** Michał Januszewski, Jörgen Kornfeld, Peter H. Li, Art Pope, Tim Blakely, Larry Lindsey, Jeremy Maitin-Shepard, Mike Tyka, Winfried Denk, Viren Jain

## Abstract

Reconstruction of neural circuits from volume electron microscopy data requires the tracing of complete cells including all their neurites. Automated approaches have been developed to perform the tracing, but without costly human proofreading their error rates are too high to obtain reliable circuit diagrams. We present a method for automated segmentation that, like the majority of previous efforts, employs convolutional neural networks, but contains in addition a recurrent pathway that allows the iterative optimization and extension of the reconstructed shape of individual neural processes. We used this technique, which we call flood-filling networks, to trace neurons in a data set obtained by serial block-face electron microscopy from a male zebra finch brain. Our method achieved a mean error-free neurite path length of 1.1 mm, an order of magnitude better than previously published approaches applied to the same dataset. Only 4 mergers were observed in a neurite test set of 97 mm path length.

## Introduction

Computers have been employed for the reconstruction of neural ‘wires’ since the 1970s ^1^mainly to capture and display the annotation decisions made by human tracers ^2–4^. The use of computers to make those decisions based on algorithms designed to detect the boundaries of cells began in earnest after new and improved approaches to the acquisition of volume EM data ^5^ started producing datasets for which the complete analysis by human annotation would be prohibitively expensive. It quickly became clear that machine learning approaches, now mostly based on convolutional neural networks (CNNs), are the method of choice ^6^. While those algorithms find most cell boundaries, the remaining error rates required the use of human proofreading, which could either take the form of inspecting the fragments (supervoxels) of an oversegmentation generated by the algorithm ^7^or by creating skeleton tracings, which then are used to gather the fragments belonging to the same cell ^8–10^.

Unfortunately, human proofreading is prohibitively expensive for larger datasets. The estimated human labor required to reconstruct a 100^3^ μm^3^ volume exceeds 100,000 hours, even when using an optimized pipeline that combines automated neural network inference and manual skeletonization ^11^. Current manual annotation workflows could be made more efficient still but are ultimately limited by the need to view all the data. A reduction of the proof-reading time by multiple orders of magnitude requires algorithms that are not only substantially more accurate but also “know their own limits” and provide a list of potentially erroneous segmentation decisions. Humans would then have to inspect only those locations ^12^.

State-of-the-art automated neurite reconstruction generally is performed in two stages. First a convolutional network infers for each image location the likelihood of there being a boundary, using the intensities of the voxels at and near that location ^13,14^. A separate algorithm (e.g., watershed, connected components, or a graph cut approach) subsequently uses the boundary map to cluster all non-boundary voxels into distinct segments ^11,15–18^. We merged the two steps by adding to the boundary classifier a second input channel, which carries the predicted object map, leading to a recurrent model. Why is this helpful? While the cost function we used to train the network is still based purely on a voxel-wise comparison with the training set, the network can now learn to make use of the fact that certain voxels in its field of view (FoV) have already been classified with high certainty in an earlier iteration. Classification decisions can be based on, for example, whether those predictions result in a more or less plausible shape for the neural process. Shape information is thus incorporated in as much as it helps the network to improve its ability to predict the boundary map. Note that this is conceptually quite different from other approaches that use cost functions that depend directly on segmentation performance ^19,20^. Here we present a realization of this concept, which we call Flood-Filling Networks (FFNs).

FFNs are trained to distinguish a single object (the foreground) from all other objects (the background). In contrast to prior machine learning-based segmentation methods ^11,16,21–23^, FFNs segment one object at a time. With each iteration the feedback pathway carries past segmentation decisions forward in time and spreads them in space. This enables the FFN, which has a relatively small direct FoV, to integrate information from far beyond its direct FoV. It also transforms the problem of single-shot classification into classification conditioned on prior predictions (i.e., classification results from nearby locations), which we believe to be a simpler problem (see Supplementary for experimental data confirming this intuition). This makes it possible for results in “easy”, unambiguous areas to inform segmentation in “hard”, more ambiguous regions.

In the following, we describe the core FFN algorithm, how FFNs can be used to reconstruct a large-scale EM volume, compare accuracy to previously published alternatives, and analyze the errors of the FFN reconstruction in detail. The quality of the fully automated segmentation obtained with FFNs is shown to far exceed previous approaches, and opens up the possibility of efficient analysis of volumes that have so far been intractable due to their size.

## Results

### Flood-filling network architecture, inference, and training

An FFN has two input channels, one for the 3D image data and another one for the current state of the “predicted object map” (POM). Using values between 0 and 1, the POM encodes for each voxel the algorithm’s estimate of whether the voxel belongs to the object currently being segmented. At each iteration of the network’s recurrent dynamics the POM is updated for all voxels in the network’s current FoV.

We generated single-voxel seeds at locations well away from the cell’s putative boundaries as detected by a simple edge filter, because we had observed that seeds placed near cell boundaries often cause neighboring objects to erroneously merge (Fig. 1a). When starting segmentation of a new object, the networks FoV is centered on the seed and the seed location’s POMs value is set to 0.95, with all other voxels set to 0.05. The values are offset from 1 and 0 to prevent the network from learning large internal weights and overfitting ^24^.

**Figure 1.**
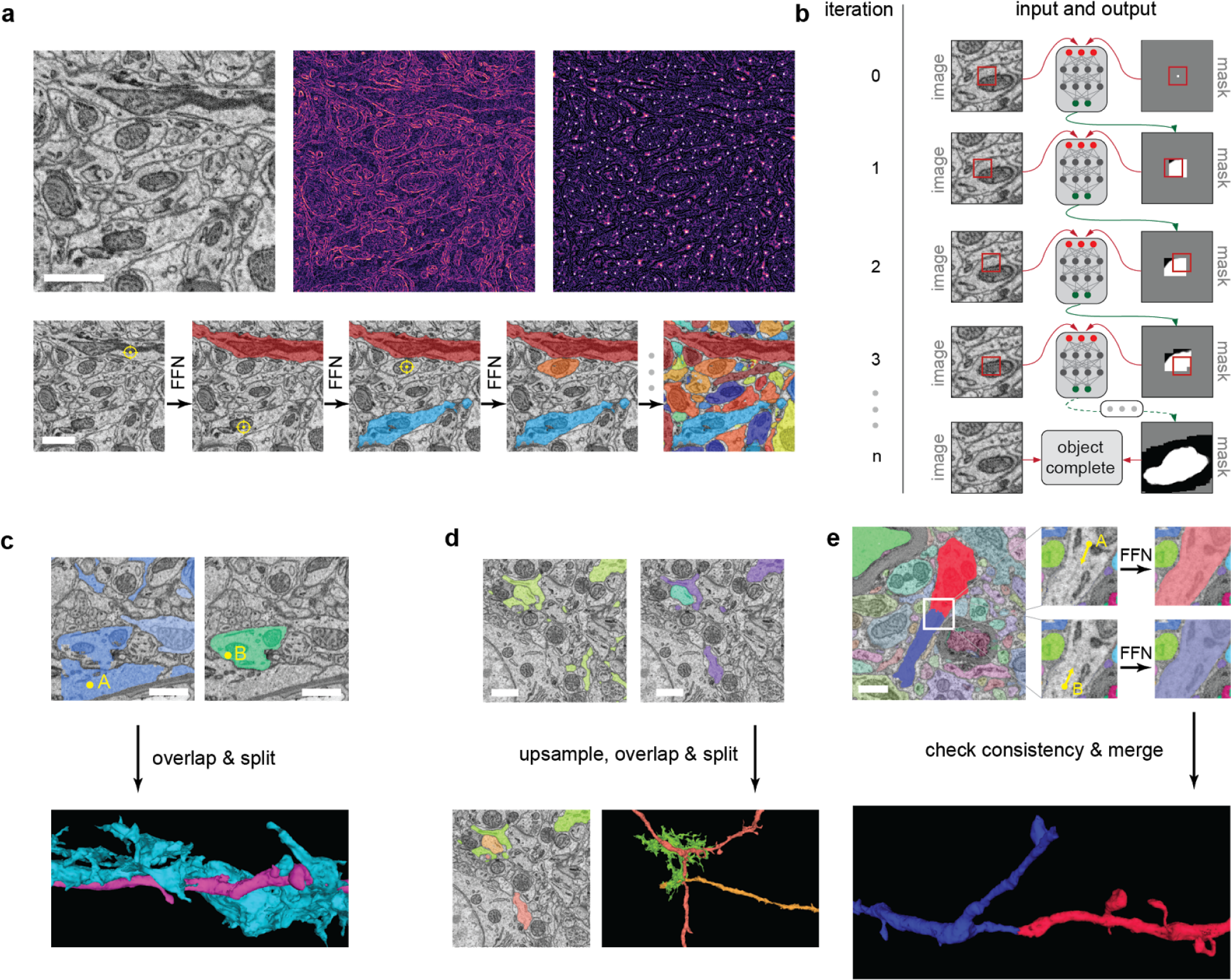
The segmentation pipeline. (a) Segmentation of a subvolume with an FFN. Top row (left to right): EM image data, local intensity gradient magnitude estimated with the Sobel-Feldman operator, Euclidean distance transform of the gradient magnitude with local peaks highlighted with white dots. The peaks are used as seed points for FFN inference. Bottom row: sequential segmentation of the subvolume with an FFN. The yellow cross-hair symbol indicates the seed point. (b) The flood-filling inference process for a single object. The red square indicates the location of the FoV in the EM data (left column) and the POM. Red lines with arrows indicate the flow of information to the inputs. Each iteration (row) consists of one forward pass of a convolutional network that receives as input both the image and the current state of the predicted object mask (POM). In the first row, the initialization of the POM is shown as specified by a single pixel (white square) within the network’s FoV (red square). Successive rows show successive iterations of FFN inference that incrementally contribute inference values to the POM while the network’s FoV moves throughout the image space. (c) Multi-seed consensus procedure. Top row: cross section through the data with the FFN segment seeded by A (left) and seeded by B (right). Note the merger between a glial fragment and a dendritic branch in the left panel. Bottom row: surface renderings of the segmentation after oversegmentation-consensus. (d) Multi-scale oversegmentation-consensus. Top row: segmentation from full resolution data (left) contains a merger between an axon and glial fragment, and the segmentation from data downsampled 2x in-plane (right) contains a merger between the same glial fragment and a dendritic branch. Bottom row: segmentation and surface rendering of multi-scale oversegmentation-consensus results in which both mergers are fixed. (e) Flood-filling agglomeration. Top row: (left) a split dendritic branch; the white square shows area of zoomed insets (right) in which FFN segmentation is started from points A (top) and B (bottom) sequentially. Bottom row: rendering of containing objects of points A and B, which are agglomerated due to satisfying mutual consistency criteria of FFN agglomeration (see text for details). The scale bars correspond to 1 μm.

The direct FoV (i.e those voxels that are directly affect the current FFN inference step) is relatively small (33×33×17 voxels, or 297 × 297 × 340 nm). Based on the inference results, the FoV may move to a new location or, alternatively, inference may terminate and generate a fixed segment from the POM (see Methods for details). Once a segment is completely fixed all seeds overlapping this segment are discarded and segmentation starts anew at one of the remaining seeds until none remain.

We applied FFNs to a 96×98×114 μm (10624 × 10880 × 5700 voxels) sized region of zebra finch brain that had been imaged with Serial Block-face Electron Microscopy ^25^ at a resolution of 9 × 9 × 20 nm. A small fraction (0.02%, 148M voxels, contained in 35 subvolumes of varying size that were distributed throughout the volume) of the data set was manually segmented by human annotators and used as ground truth to train the FFN.

At the core of the FFN is a multi-layer CNN. During each iteration step the POM values for multiple voxels are updated, in our case all voxels of the current FoV. During training the POM is first initialized by seeding a single-voxel in the center of the 49×49×25 voxel training example. A single iteration is then executed and its result is used to update the POM (Fig. 1b). The network weights are then adjusted via stochastic gradient descent using a per-voxel cross-entropy (logistic) loss ^24^. This procedure is repeated with the FoV position at a number of locations offset from the center by +/− 8 voxels (72 nm) laterally and +/− 4 voxels (80 nm) in **z** direction. In order to optimize the training procedure and remain consistent with the inference procedure, a FoV position was used only if the POM value of the new center voxel exceeded 0.9 immediately before the move. The order of moves was randomized.

### Irregularity detection and automated tissue classification to prevent segmentation errors

Many of the FFN’s errors occur at data irregularities, such as cutting artifacts or alignment mistakes. While frequent enough to affect the overall error rate, such irregularities are too rare (affecting fewer than one voxel in a hundred) for the network to learn how to avoid them. Rather than enriching them in the training set, we decided to instead detect such irregularities in a separate process. When an irregularity was found we partitioned the neural process. While such partitions were mostly splits (errors in which two processes are erroneously disconnected from one another), many of those splits were later corrected at the agglomeration stage. Objects that were not reconnected in this way could be candidates for later human proofreading.

We also observed that the segmentation quality often declined near objects, such as somata or blood vessels, that were significantly larger than the cross sections of typical axons, dendrites and the FFN’s FoV. To address this we trained a separate CNN to detect blood vessel, cell body, neuropil, myelin, or out of bounds voxels and used these classifications to prevent the FFN from extending objects beyond the neuropil.

### Hysteresis and approximate scale invariance in FFNs

FFN segmentation results were dependent on the placement of the initial seed and on the order in which objects were segmented. There were, for example, cases where a merger (two or more processes are erroneously connected to one another) was only created between two objects (A and B) when the segmentation order was A-B but not when it was B-A (“unidirectional mergers”).

This can be exploited to eliminate mergers while increasing the number of splits by only a small amount. Specifically, we compared a forward segmentation of the data with its backward segmentation, for which the the order of seeds was reversed and accepted all splits as real (which we call the oversegmentation-consensus), which means that only those mergers remained that occurred in both segmentations (Fig. 1c). Just as we used different initial seeds to build consensus analyses, we were similarly able to use analyses carried out on subsampled raw images, despite the FFNs not being explicitly trained to do so. Bidirectional mergers at full resolution often disappear completely or become unidirectional when inference is run at a reduced resolution, which we confirmed for data that were downsampled in-plane by factors of 2 or 4 and two-fold in the axial direction. Note that downsampling increases the FoV in physical space in these cases to, respectively, 594 × 594 × 340 and 1188 × 1188 × 680 nm.

We finally generated an oversegmentation-consensus using forward and reverse segmentations at the original and at down-sampled resolutions (see Fig. 1c,d and Online Methods for details). These procedures are conceptually similar to the process of ensembling in machine learning, i.e. combining the decisions of multiple different classifiers. Others have used the averaged predictions of the same classifier applied to modified versions of the raw image ^18,26^. In our case, the classifier was also kept constant, and variance was generated by changing the initial conditions and the scale of the image. These procedures increased the number of splits by a factor of only 2, but reduced the number of mergers by a factor of 82 (see also Fig. 3b,c for path length statistics).

### FFN agglomeration

In order to reduce the number of splits, we agglomerated segments throughout the volume. Unlike previous automated agglomeration approaches, which involve training a classifier to score pairwise merge decisions ^27^ or predict a compatibility score between an agglomerated segmentation and the raw image ^28^, we instead used the FFN model itself to perform agglomeration.

To determine whether a pair of segments in spatial proximity are part of the same neurite, we extracted a small subvolume (about 1 μm^3^ in size) around the point of their closest approach. We then placed seeds in parts of the two objects inside the subvolume, at locations maximally distant from object boundaries, and performed two independent FFN inference runs, one for each of these seeds, while keeping the remaining objects fixed (Fig. 1e). If the resulting POMs overlapped to a high degree (using the intersection-over-union Jaccard index as a criterion), the objects were merged. This procedure takes advantage of the sensitivity of the FFN to the seed location and allows calibration of merge decisions by varying the threshold applied to the intersection-over-union value.

### Large-scale FFN segmentation pipeline

We combined tissue masking, FFN inference, oversegmentation-consensus, and FFN agglomeration into a three-stage pipeline that was used to segment the entire volume.

1. Alignment. The sections within the volumetric dataset were precisely registered using elastic alignment ^29^, which, compared to a translation-only alignment, reduced the number of partitions introduced by the irregularity-detection procedure.
2. Cell-body segmentation. The parts of the volume corresponding to the interiors of cells bodies were segmented by running a FFN restricted to voxels labeled as being part of a cell body by the tissue-type classification, using seeds manually placed into all 454 somas in the volume (1 human hour was required for manual annotation, but this task could also be automated with 0.97 precision and 0.96 recall ^30^). Explicit handling of cell bodies was necessary due to the dramatically different spatial scale of cell bodies (average diameter 9 μm) and neuropil, which comprised the majority of training data for the FFN (average diameter 0.19 μm).
3. Neuropil segmentation and agglomeration. FFN inference was restricted to voxels labeled as neuropil by the tissue classifier, and five separate segmentations were generated and reconciled with oversegmentation-consensus. FFN agglomeration was used to merge neurites in the segmentation, which reduced the total rate of splits by 44% and increased the run length by 1149%.

### Evaluation of large-scale segmentation accuracy

The most popular datasets for comparison and evaluation of computational methods for EM reconstruction are relatively small subvolumes of tissue that have been completely segmented by hand, i.e. each pixel in the subvolume has been assigned to an object ^18,31^. First we evaluated FFN and comparison methods on a 5 × 5 × 5 μm subvolume but found that the accuracy of the methods on this subvolume did not predict a method’s performance on the 5000-times larger complete volume (for example, the fraction of ground truth neurite annotations containing a merger in the CNN+GALA baseline segmentation method increased from 7.3% to 46.3%; see supplementary for details).

Thus in order to measure the accuracy of segmentation results over length scales comparable to the path length of neurons in the complete volume, we “skeletonized” individual neurons ^8^. Human annotators used KNOSSOS software (https://knossostool.org/) to manually annotate individual neuron structure as a set of nodes and edges forming an undirected tree. We created two non-overlapping sets of skeletons. The *tuning set*, which was used to optimize the hyper-parameters of the segmentation pipelines, contained 13.5 mm total neurite path length (of which 27% was axonal) distributed among 12 neurites with a 0.8 mm median path length. The *test set*, which was used solely for evaluation purposes, contained 97 mm total path length (34% axonal) across 50 neurons (2 mm median path length). We found that the skeletons contained eight mergers and 66 splits (mostly missed dendritic spines), even after human consensus generation based on multiple independent tracings. We corrected this by having two human experts (M.J., J.K.) examine every putative merger in the automated segmentations, and then fix the manual skeletons when they were deemed erroneous.

Based on overlap with the automated segmentation, each edge of the ground truth skeletons was classified as either correctly reconstructed in the segmentation, omitted, split, or part of a merged segment^11,32^. For example, if a segment within the automated reconstruction overlapped nodes from two different skeletons, then all edges overlapping that segment were counted as “merged.”

Only about 1.4% of the total path length in the imaged volume has been manually skeletonized. This allowed us to automatically detect all splits that occurred in the skeletonized neurites, but only those mergers that were with with other skeletonized neurites. This severely underestimates the number of mergers per cell. To correctly estimate the merger rate, we also (in addition to all automatically produced segments that contained nodes from more than one skeleton) classified a segment as containing a merger if it contained at least two skeleton nodes *and* if any part of the segment extended farther than 2.2 μm away from the skeleton to which that node belonged.

Finally, we calculated an “expected run length” (ERL) that measures the average error-free path length of neurites in the automated reconstruction. Even a single omission or split terminates the accumulation of run length; segments containing mergers do not contribute any run length at all. Note that such severe penalization of mergers is useful when optimizing an automated segmentation method since it reflects the difficulty involved in manually finding and correcting mergers when proofreading. However, when evaluating a proofread segmentation, which when combined with synaptic annotations ^30^ will yield the connections matrix, a different metric that more equally weights split and mergers will reflect the usefulness of the segmentation better. See Supplementary for full details of manual skeletonization, edge classifications, and run length calculation.

Our final reconstruction (FFN-c in Fig. 3) reached an ERL of 1.1 mm and contained only 4 mergers (Fig. 2). None of these mergers were between the ground truth skeletons, and we detected them with the heuristic procedure described above.

**Figure 2.**
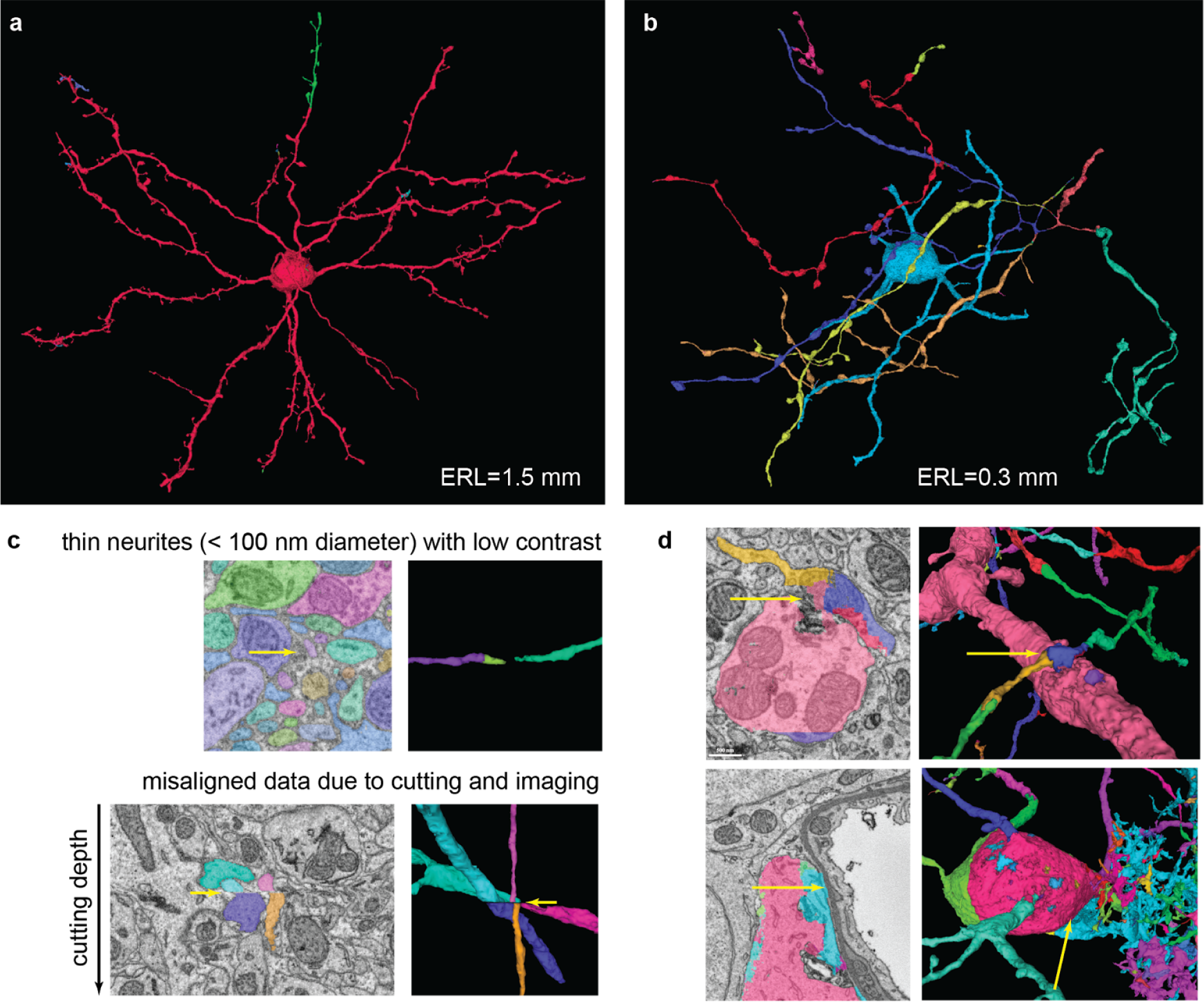
Qualitative analysis of segmentation accuracy. Different colors indicate different segments. Neurons reconstructed with the full pipeline (FFN-c) with (a) the largest (1.5 mm) and (b) shortest (0.3 mm) run lengths. (c) Zoomed views of splits caused by low contrast (top) and a slice misalignment (bottom). (d) Both rows show a set of pre-agglomeration segments (segments in FFN-b) that were erroneously merged during FFN agglomeration (segments in FFN-c). Top: dendrite-axon merger caused by small spillout of the dendrite segmentation. Bottom: cell body-glia merger caused by inaccurate cell body segmentation.

**Figure 3.**
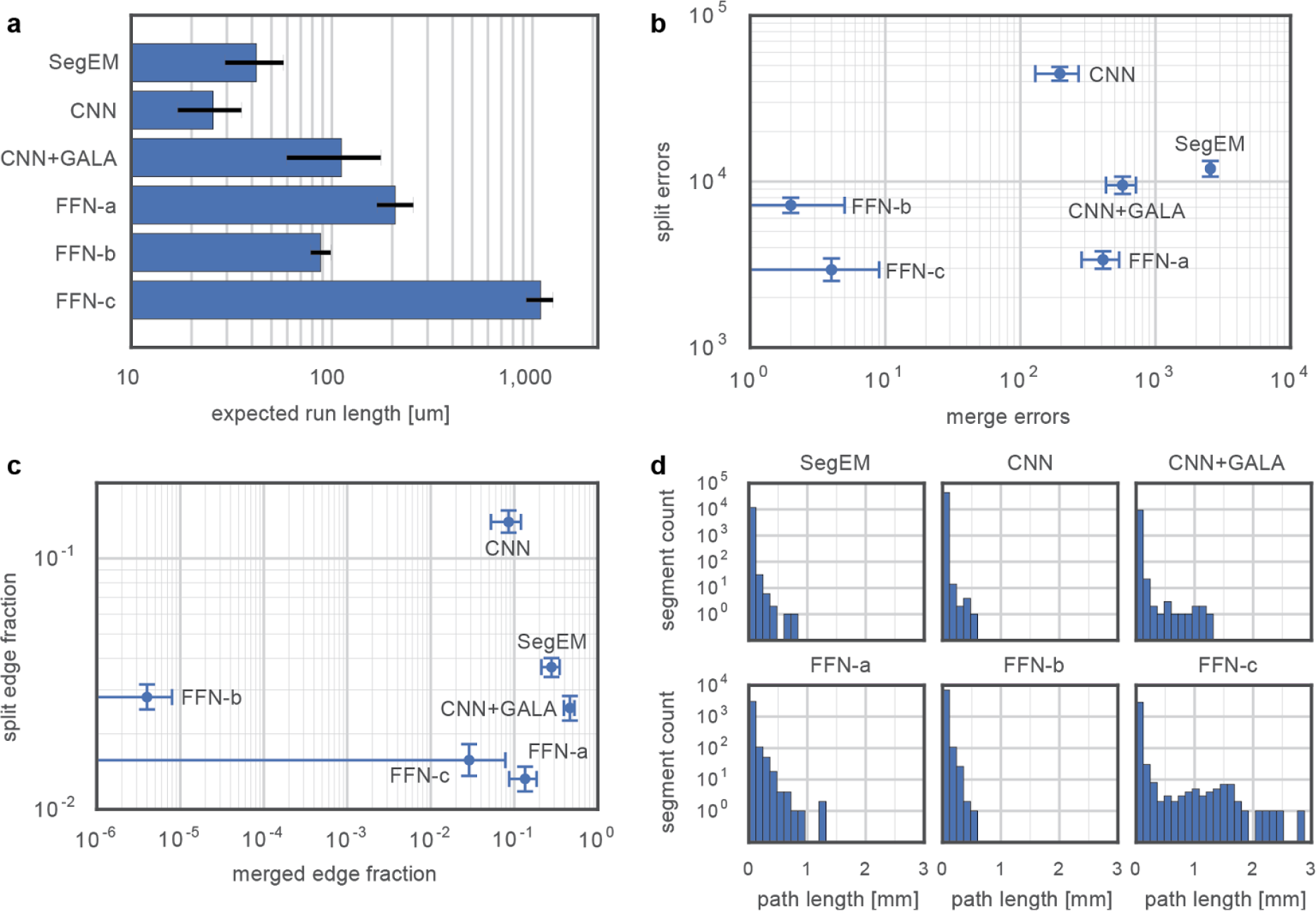
Segmentation method accuracies measured by comparison with 50 manually traced and verified skeletons (97 mm path length). SegEM: seeded watershed applied to convolutional neural network (CNN) boundary prediction ^11^, CNN: watershed over affinity graph predicted by a CNN, CNN+GALA: affinity graph watershed output agglomerated with a random forest classifier ^27^, FNN-a: single-pass FFN segmentation, FFN-b: FFN segmentation after multi-seed and multi-resolution consensus (note that the ERL decreased, but the merge rate is also decreased to almost 0), FFN-c: result of the entire FFN pipeline, including FFN agglomeration. (a) Expected run length. The end of the x-axis scale (at 2.1 mm) indicates the maximum ERL attainable for this dataset and set of ground truth skeletons. (b) Merger and split counts ^11^. (c) Fraction of ground truth skeleton edges classified as either split or merged. (d) Merge-free segment length distributions for the different segmentations. Error bars represent 95% confidence intervals and were calculated using the bootstrap method with 10,000 resamples.

To compare the performance of FFNs to alternative approaches to connectomic reconstruction, we implemented two of those approaches. The first approach, which we refer to as the “baseline,” combines a 3d convolutional neural network trained to produce long-range affinity graphs ^21^, affinity graph watershed segmentation ^33^, and random-forest object agglomeration using “GALA” ^17,27^. The convolutional network used an input FoV of 35×35×9 voxels and was trained to produce a long-range affinity graph that for every voxel predicted its binary connectivity to all its neighbors within a 3×3×1 radius. We also evaluated a recursive boundary prediction network ^30^ with a larger 201×201×21 FoV (for details of all network architectures see Supplementary), but found that it performed worse than the baseline network. The parameters for the affinity-graph watershed procedure were optimized by grid search, and GALA agglomeration was performed with a random forest classifier trained on the subvolumes of labeled data. Finally, we evaluated “SegEM” ^11^, in which 3d convolutional neural network boundary predictions are over-segmented with watershed. The same 3d convolutional network as the baseline was used, and we optimized the parameters by grid search over full volume segmentations. Among these alternative approaches, the baseline method achieved the highest ERL (112 microns; see Fig. 3 and Supplementary Table 5).

### Analysis of errors by neurite type

We manually classified fragments of neurites in ground truth skeletons as axons or dendrites, and annotated the locations of the base and the head of 182 dendritic spines. We then used this data to measure error rates of the FFN-c segmentation for the different neurite categories (see Supplementary for details). We observed that the automated reconstruction is better than human annotators in identifying dendritic spines (95% and 91% recall, respectively). While precision remained close to 100% and was slightly higher for the automated results (99.7%, 100% automated vs 98%, 99% human-generated for dendrites and axons, respectively), recall for both axons and dendrites was still inferior to human performance (68%, 48% automated vs 89%, 85% human-generated for dendrites and axons, respectively).

Many of the remaining splits could be attributed to data artifacts, which affects all types of neurites, or low contrast (Fig. 2c), which affects only low-diameter processes such as axons. We attempted to correct for misalignment and single-slice artifacts in the agglomeration procedure (see Methods), but there remain difficult cases that, while unambiguous to humans, require further improvement in automated techniques.

## Discussion

Flood-filling networks differ from other machine learning-based segmentation approaches in several ways: a recurrent network architecture, the direct generation of segments (rather than having to rely on a separate clustering step), and an inference procedure that segments objects one at a time. We also exploited several additional capabilities of FFNs, such as the ability to reduce mergers by ensembling multiple segmentations generated by varying seed point location. Applying these techniques to a roughly 1 million cubic micron volume of songbird tissue yielded automatically segmented neurons with an average error-free run length of 1.1 mm.

The main disadvantage of FFNs is the high computational cost. For example, performing a single pass of the fully-convolutional FFN network over the whole volume is 14.4x more expensive than the more traditional 3d convolution-pooling architecture in the baseline. This is because multiple and partially overlapping inference computations are required to segment both a single object as well as implement the sequential nature of multiple object segmentation. On average, every voxel of the volume was processed by the FFN 59 times in a segmentation run. Ensembling segmentations from multiple seed points further multiplies FFN inference cost by 2.38×, and agglomeration introduces another factor of 2.16×. In total, the FFN pipeline required 14.4 × 2.38 × 2.16 = 74 greater computation compared to the baseline CNN (see Supplementary for details).

However, the benefits of FFN segmentation are likely to outweigh the increase in computational cost, given the vast saving of human proofreading time that follows from order-of-magnitude improvements in reconstruction accuracy, as well as the continuously decreasing cost of computational power and potential for algorithmically derived improvements in efficiency ^34^. The very low rate of mergers in the FFN reconstruction would greatly reduce the need for manually splitting of undersegmented 3d objects, one of the most costly parts of proofreading. The large size of the automatically generated segments should make the shape-based prediction of potential splits more reliable and thus make a replacement of the laborious manual segmentation step by focussed annotation feasible. As the error-free path length increases it becomes more and more likely that a segment that contains an error also violates one of the known topological properties common to all neurites, such as the expectation that all neurites either connect to a cell body or extend to the border of (and thus beyond) the imaged volume, and such violations could become an efficient way to guide the proofreading process and to provide a measure for the residual error rate in the segmentation.

## Acknowledgements

We thank Tom Dean and Blaise Agüera y Arcas for useful discussions and support. We also thank Michale Fee and Jon Shlens for comments on the manuscript.

## Online methods

### Tissue irregularity detection

We used cross-correlation in order to detect irregularities in the input EM data caused by artifacts in the image acquisition process or by imprecise alignment of the images. For every pair of neighboring sections, 160 × 160 pixel patches were extracted, centered at every node of a 2d grid with step size 40 pixels. We then computed the normalized cross-correlation of the two patches corresponding to every grid node using FFT convolution in FULL mode (i.e., convolution results were computed at every point of overlap, even if partial, by padding with zeros). The peak in the correlation image was identified, and its offset from the center of the image was taken as an estimate of the local lateral section-to-section motion, forming a sparse 2d vector field over the whole volume. Based on experiments with FFN inference in areas affected by data irregularities, we recorded an irregularity when either component of the vector field exceeded a value of 4.

### Tissue type classifier

We trained a convolutional network to predict whether a voxel belonged to one of six categories that represented general structural features of the image volume. First, we manually labeled 26.7 million voxels (0.016% of the volume) at 2x reduced lateral resolution as either blood vessel (4.4M voxels), cell body (11.5M voxels), myelin (1.5M voxels), neuropil (7.4M voxels), or an “out-of-bounds” (1.8M voxels) category defined for those voxels in the embedding substrate that were outside the bounds of the actual songbird tissue. Manual annotation were sparsely created on every 500th slice by two authors (V. J., M.J.) using a custom web-based tool (“Armitage”) that enabled manual painting of voxels with a modifiable brush size (see Fig. 4); in total, annotation required 5 hours of human time.

We then used TensorFlow ^35^ to train a 3d convolutional network to classify a 65x65x65 patch centered on each manually labeled voxel. The network contained three “convolution-pooling” modules ^36^ consisting of convolution (3×3×3 kernel size, 64 feature maps, VALID mode where convolution results are only computed where the image and filter overlap completely) and max pooling (2×2×2 kernel size, 2×2×2 stride, VALID mode), followed by one additional convolution (3x3x3 kernel size, 16 feature maps, VALID), a fully connected layer (512 nodes, expressed as a point-wise convolution), and a six-class softmax output layer ^24^. We trained the network by stochastic gradient descent with a minibatch size of 16 and 4 replicas ^37^. During training, each of the six classes was sampled equally often. Training was terminated after 1 million updates.

Inference with the trained network was applied to all voxels in the image volume using dilated convolutions, which is several orders of magnitude more efficient than a naive sliding-window inference strategy ^38^. Finally, the analog [0,1]-valued network predictions were thresholded and used to prevent certain image regions from being segmented, as detailed in section “Large-scale FFN segmentation pipeline”.

**Figure 4.**
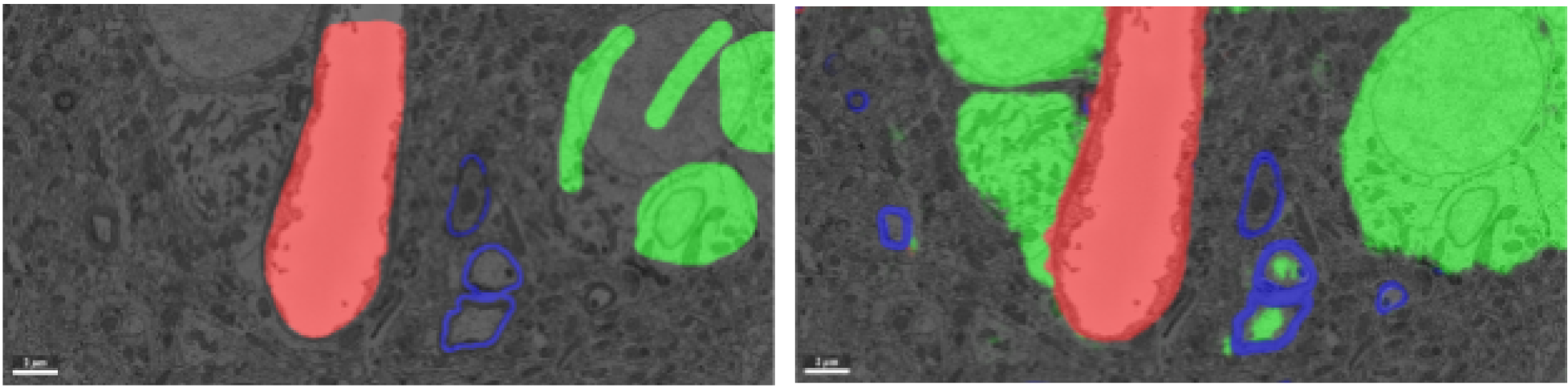
Manual annotations (left) and convolutional network inference (right) of a subset of the labeled voxel classes: blood vessel (red), myelin (blue), and cell body (green). False positive identifications of cell body voxels are visible in the automated inference (inside the myelinated area).

### Flood-filling Networks (Architecture, Training, Inference)

#### Architecture

The FFN comprised a stack of 3d convolutions in SAME mode (input and output of every layer of equal size, with input implicitly padded with zeros to achieve this) with skip connections, rectified linear (ReLU) nonlinearities ^24^, 3x3x3 kernel sizes, and 32 feature maps in every layer but the the last layer. The network consisted of 19 convolutional layers containing a total of 472,353 trainable weights (see Fig. 5). The input module consisted of a sequence of a 3d convolution, ReLU nonlinearity, and another 3d convolution. This was followed by eight residual modules that performed a ReLU nonlinearity, 3d convolution, ReLU nonlinearity, and 3d convolution. The last layer performed a voxel-wise convolution that combined input from all feature maps (1×1×1 kernel size with a single output feature map). The input and output of the network were equal in spatial size -- 33×33×17 voxels. The input was formed by a 2-channel image, containing EM data in channel 1 (normalized to 0 mean and unit standard deviation) and the current state of the predicted object mask in logit form in channel 2. The output of the network was the updated state of the predicted object mask in logit form.

**Figure 5.**
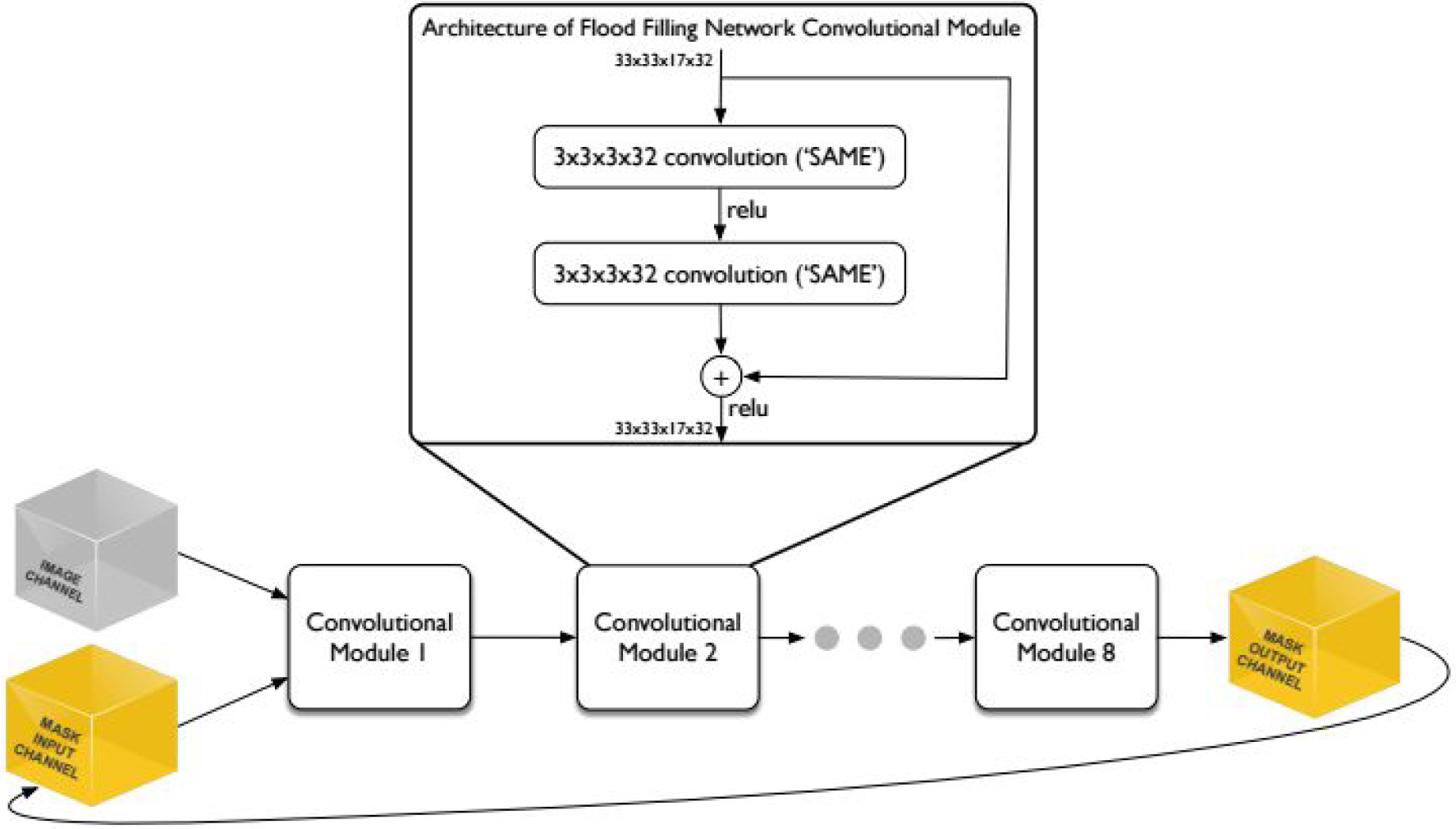
Architecture of the FFN. The internal modular architecture used here corresponds to “full pre-activation residual modules” ^39^. The architecture was chosen because in our experiments it showed better convergence than alternatives (no skip connections, or other proposed variants of skip connections ^40^). We note that our network contains significantly fewer weights than those used in many recent works (e.g. 74x times fewer than ^18^).

The FFN was implemented in TensorFlow ^35^ and trained with voxelwise cross-entropy loss:

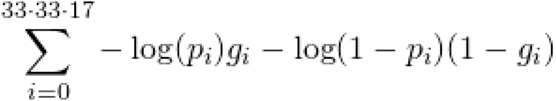

where p_i_ is the predicted voxel value and g_i_ is the ground truth label after smoothing. Training proceeded for 7 days using asynchronous stochastic gradient descent at a learning rate of 0.001, in a distributed setting with 32 NVIDIA Tesla K40 GPUs and batches of 4 examples.

#### Training example sampling

The initial set of training examples was formed by extracting all subvolumes 49 × 49 × 25 voxel in size and fully contained within one of the 33 regions densely segmented by human annotators. The size of the subvolume was chosen to allow FoV movement by one 8-voxel step in every direction.

The ground truth segmentation within every subvolume was binarized by setting voxels belonging to the same object as that of the central voxel of the subvolume to 0.95, and the rest of the voxels to 0.05. These soft labels ^24^ provided the desired object mask probability map that the FFN was trained to predict.

For every one of the initial training examples, the fraction (f_a_) of active mask voxels was a calculated. The training examples were then partitioned into 17 classes, such that an example was assigned to class *i* if *t*_*i*−1_ ≤ *f*_*a*_ < *t*_*i*_, and *t= (0.0, 0.01, 0.02, 0.03, 0.04, 0.05, 0.06, 0.075, 0.1, 0.2, 0.3, 0.4, 0.5, 0.6, 0.7, 0.8, 0.9, 1)*. For example, a training example with *f*_*a*_ = 0.5 would be a assigned to class 12. During training, each of the 17 classes was sampled equally often.

#### Seed selection

The seed points for FFN inference were selected as follows: all pixels where the 3d Sobel-filtered EM image was larger than the same image filtered with a Gaussian with σ = 49/6 was set to 1, and 0 otherwise. We then computed the Euclidean distance transform of the resulting binary image and selected local peaks of that transform as the initial FFN seeds.

These seeds were then consumed serially in raster order. All seeds found to be within 3 voxels or less from an existing segment at the time of the inference start were discarded.

#### Field of View movement procedure

The FoV of the FFN was moved using the following procedure. A list (Q) of positions to be visited was initialized with a location obtained from the seed policy. In the segmentation loop, a location (x, y, z) was extracted from the head of Q, and the FoV was moved to that position, which was then marked as visited. Visited locations were stored in order to ensure that every location was visited at most once during segmentation of an object; locations were stored at 8x reduced resolution in the XY directions and 4x reduced resolution in the Z direction. This reduced resolution effectively determines the minimum step size by which the FoV can be moved, and was used to control the efficiency of inference.

After an inference call, all POM voxels within the FoV were updated, except those that were previously updated by the FFN, had a prior value < 0.5, and a new value larger than the prior value (this biased the network towards splits in areas where predictions of background/foreground were not consistent between iterations). A cuboid of POM values (x-Δx <= x <= x + Δx) ⸀ (y-Δy <= y <= y + Δy) ⸀ (z-Δz <= z <= z + Δz) was then extracted, and the maximum value was then identified on every one of its faces. Whenever this value matched or exceeded the movement threshold of 0.9, the corresponding location was appended to Q unless it was visited before.

The inference loop was terminated when Q was empty. At that point, if the number of voxels with POM values >= 0.6 was >= 1000, a new segment was created consisting of those voxels, otherwise the POM values were reset to 0.05 without creating a segment. Segmentation was terminated when no more seeds were available to start new inference runs. We used Δx = Δy = 8, Δz = 4 which were the largest values that did not result in an increased number of errors in our tests, while remaining computationally tractable.

#### Criterion for model evaluation and selection

In addition to the densely labeled ground truth data, a 560x560x250-voxel region of the J0126 volume was exhaustively skeletonized by human annotators using Knossos, resulting in 221 skeleton fragments within the subvolume. We used this subvolume to optimize the FFN performance.

During training, a snapshot of the network weights (“checkpoint“) was saved every hour. After training was completed, we ran FFN inference over the densely skeletonized subvolume with every available checkpoint, and evaluated the resulting segmentation with skeleton metrics. We selected the checkpoint that had the highest expected run length among the set of checkpoints with the least number of mergers (in our case this corresponded to the set of checkpoints with zero mergers).

#### Distributed inference

In order to perform FFN inference efficiently over the whole 663 GB dataset, we split it into overlapping 500x500x500-voxel subvolumes. The subvolume corners were located on a regular grid with a step size of 436 pixels, so that neighboring subvolumes overlapped by 64 voxels in every direction. We ran FFN inference as described above for every subvolume independently, distributing the computational load over a cluster of machines that contained GPUs.

The global segmentation was built using these partial segmentations. A “core” segmentation was extracted from every subvolume by discarding a 32-voxel wide envelope (a subset of the overlap area) and computing connected components of the remaining segmentation. For every face of a subvolume *A*, a 1-voxel thick plane parallel to this face was extracted from the middle of the overlap area. A corresponding plane was extracted from the neighboring subvolume *B* sharing the given face. For every segment s_A_ in the A plane, the maximally overlapping (by number of shared voxels) segment s_B-A_ was identified, and vice-versa. Segments for which s_A-B_ = s_B-A_, i.e. which were mutually maximally overlapping, were then merged. This conservative merging procedure was used in order to avoid spurious mergers (−84%) when creating the global segmentation, at a cost of increased splits (+28%).

### Multiresolution oversegmentation consensus

The oversegmentation-consensus procedure relies on intersecting objects in voxel-space. In the case of consensus between segmentations at different resolutions, we upsampled the lower resolution segmentation with nearest neighbors interpolation. We then applied seeded watershed segmentation to the Euclidean distance transform of the higher resolution segmentation as the height map using the upsampled segmentation as seeds. This did not change the topology of the upsampled segmentation, but prevented voxel-level differences between the two segmentations from generating new segments in the oversegmentation-consensus procedure.

To reduce the number of splits in the multi-resolution oversegmentation-consensus procedure, the lower resolution segmentation was filtered by eliminating all objects containing fewer than 100,000 voxels before upsampling. Any object consisting of fewer than 1000 voxels was also removed after consensus.

### Cell body segmentation

We created a separate segmentation containing only cell bodies, and used it as a starting point for all subsequent FFN inference. To do so, we performed three FFN inference runs at resolutions of 9x9x20 nm (original), 18x18x20 nm and 36x36x40 nm, with areas of the volume not classified as a cell body by the tissue type classifier masked out. We then applied multi-resolution oversegmentation-consensus, and removed all objects with fewer than 10M voxels. The segmentation was resampled at an isotropic resolution of 160 nm. At this resolution, we computed the Euclidean distance transform within the cell body segments, and used seeded watershed with manually placed seeds (cell body center annotations) to separate adjacent cell bodies. The corrected segmentation was upsampled back to the original resolution of the dataset, and the separated cell bodies were used as seeds for a watershed transform. Background voxels of the full-resolution uncorrected cell body segmentation were masked out so that no new voxels were labeled by watershed.

### Flood-Filling Network Agglomeration

#### Candidate object pair generation

A subset of all possible supervoxel pairs were considered for automated agglomeration. Specifically, we computed agglomeration scores (see below for details) for any pair of objects where both supervoxels contained at least one voxel within the same 5x5x5 cuboid radius. For each such supervoxel pair, we also computed a “decision point” defined as the midpoint of the shortest line that connects any two points of the supervoxels.

For every such decision point involving two objects, A and B, both containing at least 10,000 voxels, we extracted a (101×101×51)-voxel subvolume of EM data and segmentation, centered at the decision point. We then removed the segments A and B from the subvolume, and ran FFN inference within it twice -- once starting from a voxel originally labeled as A, and once starting from a voxel originally labeled as B. To identify the starting voxel, we computed the Euclidean distance transform of the segment A or B, and chose the voxel with maximum distance as the FFN seed point. In case the inference run failed to generate an object covering at least 60% of the voxels of the original object (A or B), a cuboid area of radius (8, 8, 4) voxels around the seed point was zeroed-out in the distance transform, and FFN inference was attempted again from a new location selected as described above. This procedure was repeated up to 16 times.

#### Agglomeration scoring

We analyzed the FFN inference results for every candidate object pair as follows. The POMs were thresholded at 0.5 to generate a binary segmentation. We computed the number of voxels in the generated segments (N_A_, N_B_ -- starting from the original segment A, and B, respectively), the fraction of voxels of the original segments within the analysis subvolume reconstructed in the generated segment (f_AA_, f_AB_, f_BA_, f_BB_),, the Jaccard index J_AB_ between the two segments (defined as the size of the intersection of the two sets of object mask voxels divided by the size of the union of the two sets of object mask voxels), and the number of “deleted” voxels d_A_, d_B_, where a voxel in the POM was considered “deleted” if during inference it was updated by the FFN from a value >= 0.8 to a value <= 0.5.

#### Agglomeration steps

We performed three runs of FFN agglomeration. The first run covered all decision points. The second run was limited to decision points affected by tissue irregularities. Specifically, we computed the cross-correlation of neighboring (*z* + *1*) and next-neighboring (*z* + *2*) sections within the agglomeration subvolume and computed local shift vectors as described in “Tissue irregularity detection”, yielding **m_z_** and **m_z+1_**. If **m_z_** satisfied the tissue irregularity criteria, but **m_z+2_** did not, then we assumed that a single-slice imaging artifact had been identified and replaced section *z* + *1* with image data from section *z* + *2*. The subvolume was then realigned with translation-only alignment utilizing the neighboring slice shift vectors, and FFN inference was performed without shift masking. The third run was performed with a subvolume of larger size -- (201×201×101) voxels, and was limited to decision points that in the first run resulted in f_A*_ or f_B*_ < 0.4.

We then merged segments A and B if f_AA_, f_AB_, f_BA_, f_BB_ >= 0.6 (at least 60% of the voxels of both original segments were reconstructed in both runs), d_A_/N_A_ < 0.02 or d_B_/N_B_ < 0.02 (only a small fraction of the object mask voxels got “deleted” during inference, in at least one of the runs), and J_AB_ >= 0.8 (segmentations starting from A and B were mutually consistent in voxel space). We then treated the merged segments as a weighted graph, with the segments as nodes and J_AB_ as edge weights, computed the connected components of this graph, and checked if any of the components contained more than one segment associated with a cell body (a cell body-cell body merger). If it did, we identified the shortest path between two such cell body segments, and removed the edge with the lowest weight along that path. This procedure was repeated until all mergers were resolved. All parameters used above were optimized using the “validation” set of 12 ground truth skeletons.

## Supplementary

### Elastic alignment of the EM stack

The raw EM sections were first translationally aligned. Cross-correlation of neighboring sections was computed to find a globally optimal shift correction for every section. This formed the initial EM stack, which was then elastically aligned using the method of reference ^29^.

We modified the method in several ways. First, block matching locations were decoupled from the elastic mesh vertices (block matches were searched for on a 200-pixel grid and mesh with 500-pixel edges was used). Second, for each block match, a spiral of offsets around each grid location was analyzed until the identification of an acceptable match, which we found to improve results in regions with poor or ambiguous texture. Finally, a conjugate gradient solver was used to relax the mesh, which we found to be less sensitive to integration step size and spring stiffness than Euler’s method, and which resulted in overall faster convergence.

Elastic alignment used 12 million patch match correspondences (i.e., tiepoints) between adjacent sections, and an additional 12 million between pairs of sections that were separated by an intervening section.

In the translationally aligned volume provided as input to elastic alignment, the tiepoints between adjacent sections had a mean residual error of 1.8 pixels; the 95th percentile error was 3.0 pixels. For the 12 million tiepoints across non-adjacent sections, the means and 95th %ile errors were 2.8 and 6.0 pixels.

After elastic alignment, these stats were 1.6 pixels (mean) and 2.0 pixels (95th %ile) between adjacent sections, and 2.6 and 5.0 pixels between non-adjacent sections. The mean of the magnitude of the displacement of the 3.3 million nodes of the elastic meshes used to model each section was 0.6 pixels.

### Precision/recall estimation

To estimate recall and precision of a single human annotator and the automated reconstruction (FFN-c) we used manually generated and proofread skeletons after (see “Ground truth verification” below for details) and dendritic spine head/base location annotations. The skeletons were manually separated and classified as axons or dendritic branches.

Dendritic spine recall was measured by comparing the segmentation at two manually placed locations, one at the base, the other at the head of a spine. If the segmentation labels at base and head were different, a false negative was counted. A spine was counted as found by a human skeleton if a skeleton node was within 250 nm of the base and head location. A match of only one of these locations was counted as false negative, and no match ignored, which means that the entire branch was missed.

For dendrites and axons, single-annotator skeleton nodes were matched to the nodes of the ground truth skeletons within a radius of 800 nm. The matched path length was counted as true positive, the unmatched path length in the ground truth skeleton as false negative, and the unmatched path length in the single-annotator skeleton as false positive. For segmentation, we computed precision and recall for every segment overlapping the ground truth skeletons, with the path length of the fragment of the ground truth skeleton overlapping the segment as true positive, and the path length of the automatically generated skeleton of a segment detected as merged and not matched with the ground truth skeleton as false positive. We computed recall and precision as a weighted sum of the per-segment recall and precision, with a weight of (path length of the fragment of the ground truth skeleton overlapping the segment) / (total path length of the ground truth skeleton). This provides the expected value of precision/recall provided that segmentation is started from a random location on the ground truth skeleton.

### Skeleton edge accuracy classification

Segmentation quality is often evaluated with respect to ground truth pixel-wise labels or object masks ^22^, but creating such ground truth for large-scale EM datasets that span billions or trillions of voxels is highly laborious. A more efficient way to generate ground truth representations of large-scale neuron topology is to “skeletonize” neurons into a collection of points that typically constitute an undirected tree ^8^.

We propose a set of metrics to evaluate a segmentation using such ground truth skeletons. Similar to previous approaches, we classify individual edges in skeletons as *correctly* or *incorrectly* reconstructed based on the presence of mergers or splits that affect nodes attached to an edge (^11,32^). We assume:

- a ground-truth skeleton *S*_*i*_ consists of edges *{e_i_, e_2_,…, e_|S|_}*,
- an edge *e* is defined by two 3-d node coordinates *A(e)* and *B(e)*,
- *S(e)* denotes the ID of the ground-truth skeleton containing edge *e*,
- *R* denotes a predicted segmentation to be evaluated, and *R(p)* returns the value (object ID) at point *p*. *R(e)* denotes either *R(A(e))* or *R(B(e))* where this is unambiguous.

An edge *e* is defined as *correctly reconstructed* if both of its nodes belong to the same object in the reconstruction and if that object does not contain any nodes from different skeletons. More formally, we classified every edge *e* into one of four categories (see Sup. Fig. 1):

- *omitted* if *R(e)* = 0
- *split* if *R(A(e))*  *R(B(e))*,
- *merged* if there exists an edge *e*_*m*_ such that *R(e)* = *R(e_m_)* but *S(e)* ≠  *S(e_m_)*,
- *correct* if none of the above is true.

The *edge accuracy* is the percentage of correctly reconstructed edges over all the ground truth skeletons, and incorrect edges can be further subdivided into the percentage of edges which have a merge, split, or omitted errors.

The definition of a merged node assumes that there is a skeleton for every object in the volume of interest. Some mergers could remain undetected when this assumption is violated, which is the case for large volumes where it is impractical to skeletonize every object manually. To mitigate this, we apply an additional merge detection heuristic. We call a segment *T* merged if there exists a point *p* where *R(p)* = *T* and *p* is more than *2.2* μm away from any skeleton node lying within *T*. The distance threshold was chosen based on the size of the neurites in the J0126 dataset and edge lengths in the ground truth skeletons. The merge detection heuristic was not applied in the vicinity of the cell body associated with the ground truth skeleton (when present in the dataset).

**Supplementary Figure 1.**
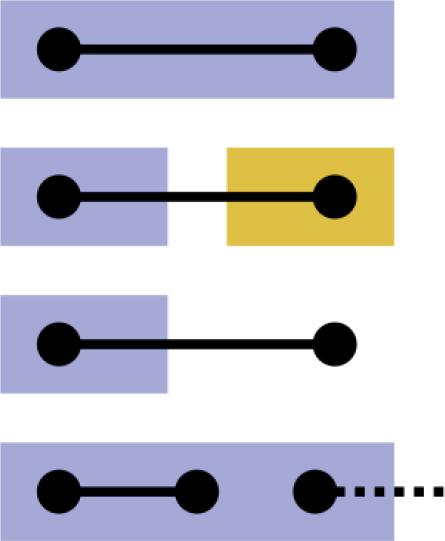
Edge classes for skeleton accuracy computation. Colors correspond to segment IDs. From top to bottom: correct edge (both nodes have the same ID), split edge (nodes assigned to different segments), omitted edge (one or two nodes do not have an associated ID), merged edge (node assigned to a segment that covers more than one skeleton).

### Expected Run Length

In order to evaluate automated segmentation results, a metric is needed that compares volumetric 3d components to ground truth skeletons and computes a single score or run length. We used the “expected run length” (ERL), which is defined as follows.

For a given skeleton S, let *CE(S)* denote the set of correct edges (as defined above). We would like to partition this set into “correctly reconstructed components” (CRCs) -- subsets of edges corresponding to valid (without a merger) segments in *R*. By definition the set *{R(e): e ∈ CE(S)}* contains only such segments.

We therefore partition *CE(S)* by the segment label *L* and define a correct component as:

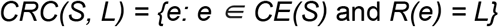

The expected run length (ERL) is the expected size of the segment that contains a randomly selected skeleton node, assuming tracing starts from a random node of the skeleton:

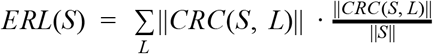

where the skeleton size is 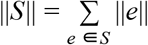

Note in particular that under this definition starting from a point which belongs to an incorrect edge (i.e. omitted, split, or merged) does not allow us to trace any correct path length and therefore does not contribute to the ERL. The ERL corresponds to the average segment length if the average is taken with the number of skeleton nodes in each segment as its weight.

The ERL for a set of skeletons *{S_k_}* is defined as:

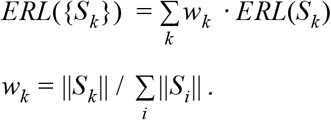

### Expected Run Length compared to other metrics

In contrast to prior approaches, the ERL takes into account the spatial distribution of errors. Previously proposed metrics, such as the total error-free path length (TEFPL) ^11,32^ and inter-error distance (IED) ^11^ are defined as simple averages and are thus insensitive to the distribution of lengths of the correctly reconstructed fragments (see Sup. Fig. 2 for an illustration).

Berning et al. define the number of splits and mergers in a predicted dense segmentation with respect to a set of ground truth skeletons as follows ^11^:

1. An individual skeleton is said to *correspond to* a predicted segment if at least *k* nodes within the skeleton are contained within the predicted segment, for *k* >= 1 or 2. Larger values of *k* provide robustness to the imprecise placement of skeleton nodes.
2. For each skeleton, the number of splits is determined as max(0, number of corresponding predicted segments-1).
3. For each predicted segment, the number of mergers is determined as max(0, number of corresponding skeletons-1).

**Supplementary Figure 2.**
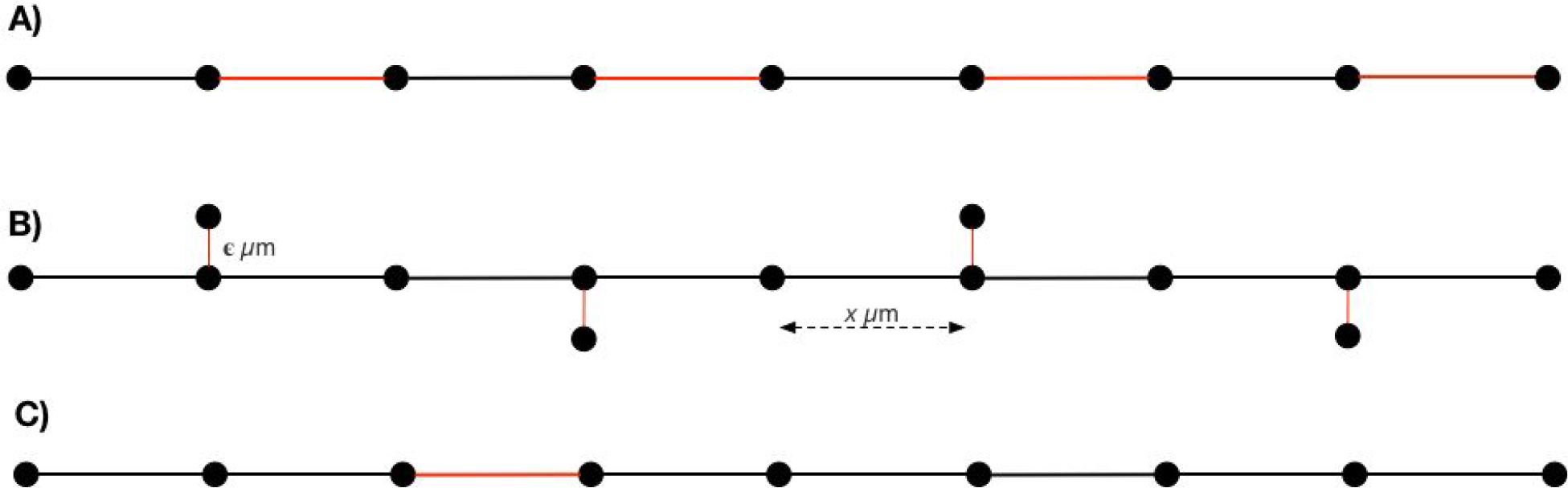
A) An idealized skeleton fragment representing *8x* μm length of neuropil where alternating edges (red) have been identified as incorrectly reconstructed in some candidate segmentation. Total error free path length (TEFPL) proposed by Pallotto et al computes an accuracy of 50% by dividing correct edge path length by total path length (4μm / 8μm), whereas the expected run length (ERL) for this fragment is *x* μm. The inter-error distance (IED) of Berning et al. is also x μm. B) An idealized skeleton fragment with “spines” of length ***ϵ*** as **ϵ**→0, the ERL and TEFPL converge to *8x*, while the IED converges to 2 μm (8 μm / 4) C) Single split dividing the process into two segments of unequal length. TEFPL is 7 μm, IED is 4 μm and ERL is 3.625 μm.

Berning et al’s definitions result in a metric with different properties from the one we propose:

- In their metric, erroneously connecting two segments can **decrease** the total number of mergers; in the limit case of a single predicted component encompassing the entire volume, the number of mergers is (number of skeletons-1).
- In their metric, the spatial distribution of splits and mergers is not taken into account: splitting off a single synapse is a single error, as is splitting a neuron in half.

In addition to being sensitive to the spatial distribution of errors, the ERL penalizes mergers very heavily since *all* edges covering the merged segments are considered erroneous, and edges marked incorrect do not contribute to expected run length. We argue that this a desired property of the metric, reflecting the significantly higher effort required to fix such errors manually during proofreading of the automated results.

### Ground truth verification

We found that the initial ground truth skeletons used for segmentation evaluation contained a substantial number of errors, despite being created by at least twofold redundant skeletonization (^8–10^) with manual resolution of discrepancies. To mitigate this, two of the authors (M.J., J.K.) inspected every discrepancy in each of the segmentations listed in Fig. 3, and when both agreed that a mistake was made in the skeletonization rather than by the FFN segmentation, the skeletons were updated. The procedure was repeated until no more error cases were detected. See Sup. Table 1 for a summary of the changes made.

Note that because of the merge detection heuristic we use, splits in the ground truth skeletons (e.g. missing neurite branches) are treated as mergers in the segmentation, and segments detected as merged do not contribute to the ERL. The quality of the ground truth data becomes increasingly more important as the path length of correctly reconstructed components improves. As an edge case, consider a complete correctly reconstructed cell with all its branches, and a single spine missing in the corresponding ground truth skeleton. This would cause the whole reconstructed object to be considered merged, and the ERL would be 0 nm.

Because of the distance threshold used in the merge heuristic (2.2 μm), a number of smaller spines missing in the skeletons remain uncorrected. The number of missing spines in Sup. Table 1 should therefore be treated as a lower bound.

**Supplementary Table 1.** Errors identified in the ground truth verification process. In total, 665 μm of skeleton path length was added (to fix splits), and 166 μm was removed (to fix mergers). Additionally, 5 skeleton nodes were moved due to their initial placement outside of the neurite represented by the skeleton.

| Fragment type | Splits<br>(untraced fragments) |  | Mergers<br>(mistraced fragments) |  |
| --- | --- | --- | --- | --- |
|  | Number of errors | Affected skeletons | Number of errors | Affected skeletons |
| dendritic spine | 59 | 13 | 0 | 0 |
| axon | 4 | 2 | 5 | 4 |
| dendrite | 2 | 1 | 1 | 1 |
| cilium | 1 | 1 | 0 | 0 |
| glia | n/a | n/a | 2 | 2 |

### Segmentation parameters

The neural network used in the baseline CNN method had the following architecture: 2d convolution (VALID mode, 5x5 filter size, 64 output features), 3d convolution (VALID mode, 5x5x5 filter size, 64 output features), 2d pooling (SAME mode, 2x2 stride, 2x2 filter size), 3d convolution (VALID mode, 5x5x5 filter size, 64 output features), 2d pooling (SAME mode, 2x2 stride, 2x2 filter size), 2d convolution (VALID mode, 5x5 filter size, 512 output features), pointwise convolution (147 output features). The output features of the last layer were treated as a 7x7x3 long-range affinity graph. The FoV of the CNN was 35x35x9. To see if a larger FoV could improve results, we also evaluated a recursive CNN ^30^ that was trained to predict a boundary map. The approach used two convolution-pooling networks: bar_b_ with a 111x111x13 FoV, and bar_br_ with a 91x91x9 FoV (see Supplementary Table 3 of ^30^ for the detailed architecture of both networks). The first network (bar_b_) took the image as input and predicted a boundary map. The second network (bar_br_) took the image and the predictions of the first network, and predicted an updated boundary map. We used the output of the bar_br_ network, which had an effective FoV of 201x201x21, and performed a grid search for watershed parameters, optimizing for ERL computed within the densely skeletonized subvolume. The best ERL found this way was 0.7 μm, which was less than the baseline CNN. We therefore decided to exclude this network from further experiments.

SegEM, as well as all FFN segmentations, used the elastically aligned volume. CNN and CNN+GALA used the original volume which was only translationally aligned, as these methods applied to the elastically aligned volume showed worse performance as measured by the set of 12 skeletons used for hyperparameter tuning. Myelin, OOB, and blood vessels were masked out in all segmentations -- voxels classified into one of these three categories were set to 0 (background).

For SegEM, the best performing set of parameters was found to be: r=0 (no morphological filtering) and h=0.045. For CNN, the optimal parameters found were T_h_ =0.945, T_l_ =0.945, T_e_=0.5, T_s_=1000). For CNN+GALA, the agglomeration threshold was set at 0.9.

### SegEM metrics

In Supplementary Table 2 we present SegEM metrics for the segmentations discussed in Fig. 3. For the SegEM segmentation method, we found the inter-error distances to be consistent with those previously reported (optimal IED = 11 μm in the present work, and 7.9 μm and 4.9 μm for the two datasets discussed in the original paper^11^).

**Supplementary Table 2.**
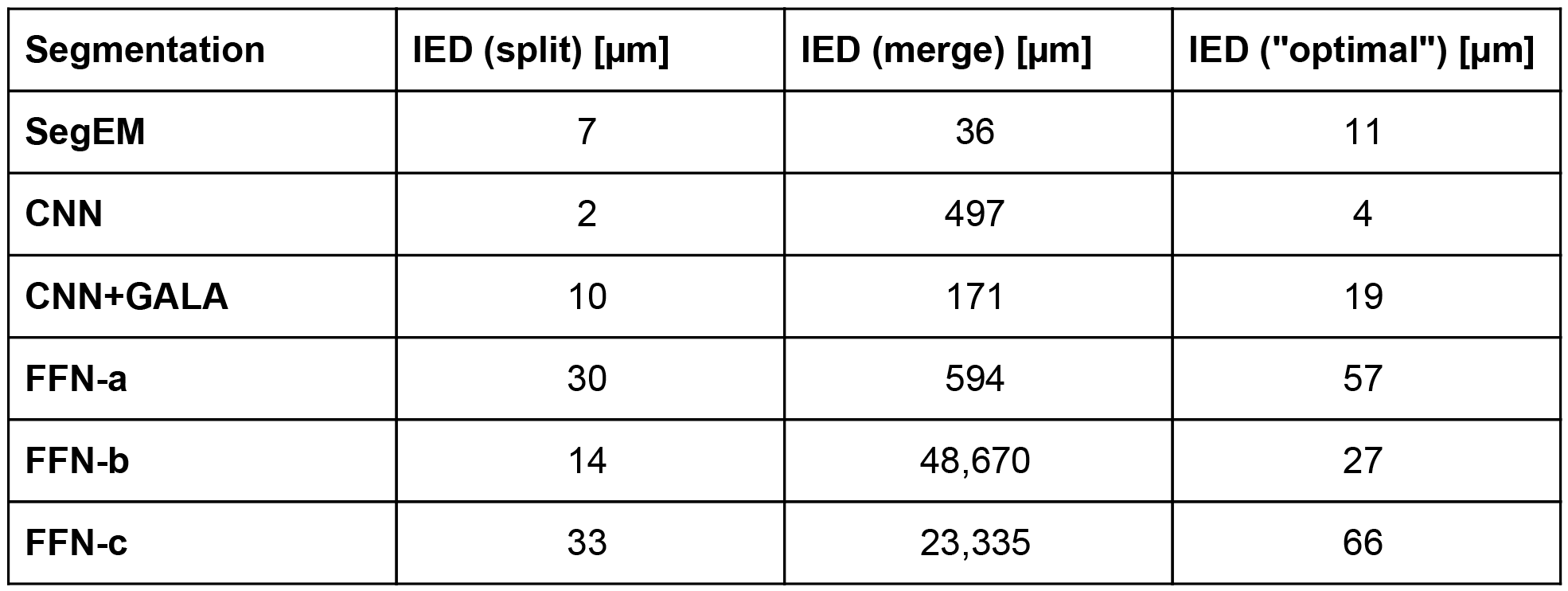
Full-volume evaluation of segmentation quality with SegEM metrics and 50 ground truth skeletons. The merge detection heuristic discussed in “Skeleton Edge Accuracy Classification” has been applied to detect mergers. Segments detected as merged were counted as a single merge error. The harmonic mean of IED split and IED merge was used to compute the optimal IED.

### Breakdown and comparison of computational cost

**Supplementary Table 3.** Computational cost of FFN inference. No pre- and post-processing of the data is taken into account in the calculations. The wall-clock time is an empirical measurement based on average inference speed with single precision floating point numbers, on a single NVIDIA Tesla P100 GPU with TensorFlow, and using CuDNN v6 as the computational backend.

| **FFN inference run** | **FFN inference calls**<br>**[x 10<sup>9</sup>]** | **EFLOPs** | **Wall time with 1000**<br>**Tesla P100 GPUs**<br>**[h]** |
| --- | --- | --- | --- |
| **9x9x20 nm, forward** | 2.22 | 41.05 | 3.15 |
| **9x9x20 nm,**<br>**backward** | 1.98 | 36.69 | 2.76 |
| **18x18x20 nm,**<br>**forward** | 0.55 | 10.18 | 0.77 |
| **18x18x20 nm,**<br>**backward** | 0.48 | 8.89 | 0.67 |
| **36x36x40 nm** | 0.05 | 1.00 | 0.09 |
| **agglomeration** | 6.10 | 112.76 | 8.64 |
| **total** | 11.39 | 210.58 | 16.08 |

For comparison of segmentation cost between the FFN pipeline and the baseline CNN in the main text, we assumed that CNN inference was done in a distributed setting with overlapping subvolumes of size 182x182x158 (the size of the subvolume was limited by the need to store the intermediate feature maps in GPU memory).

### Local versus Global Evaluations

In our experiments, we have repeatedly found that local evaluations using small (order of hundreds of μm^3^) subvolumes of data underestimate error rates. Similar observations were made in the context of synapse prediction in ^30^. In Sup. Table 3 we provide evaluations of the segmentations from Fig. 3 over the 5 μm × 5 μm × 5 μm densely skeletonized subvolume (with connected components of the segmentation recomputed after restricting it to the subvolume). The total skeletonized path length is 1 mm (20% of which is glial), and the maximum possible ERL is 13.4 μm.

Note that if two objects were merged outside of the subvolume, but were directly adjacent to each other within the subvolume, recalculation of connected components would not allow them to be split. The number of merges is therefore overestimated compared to what it would be if the segmentation procedure was restricted to the subvolume from the beginning (cf. FFN-c in Sup. Table 4 with and “recurrent single object (FFN)“ in Sup. Table 6).

**Supplementary Table 4.**
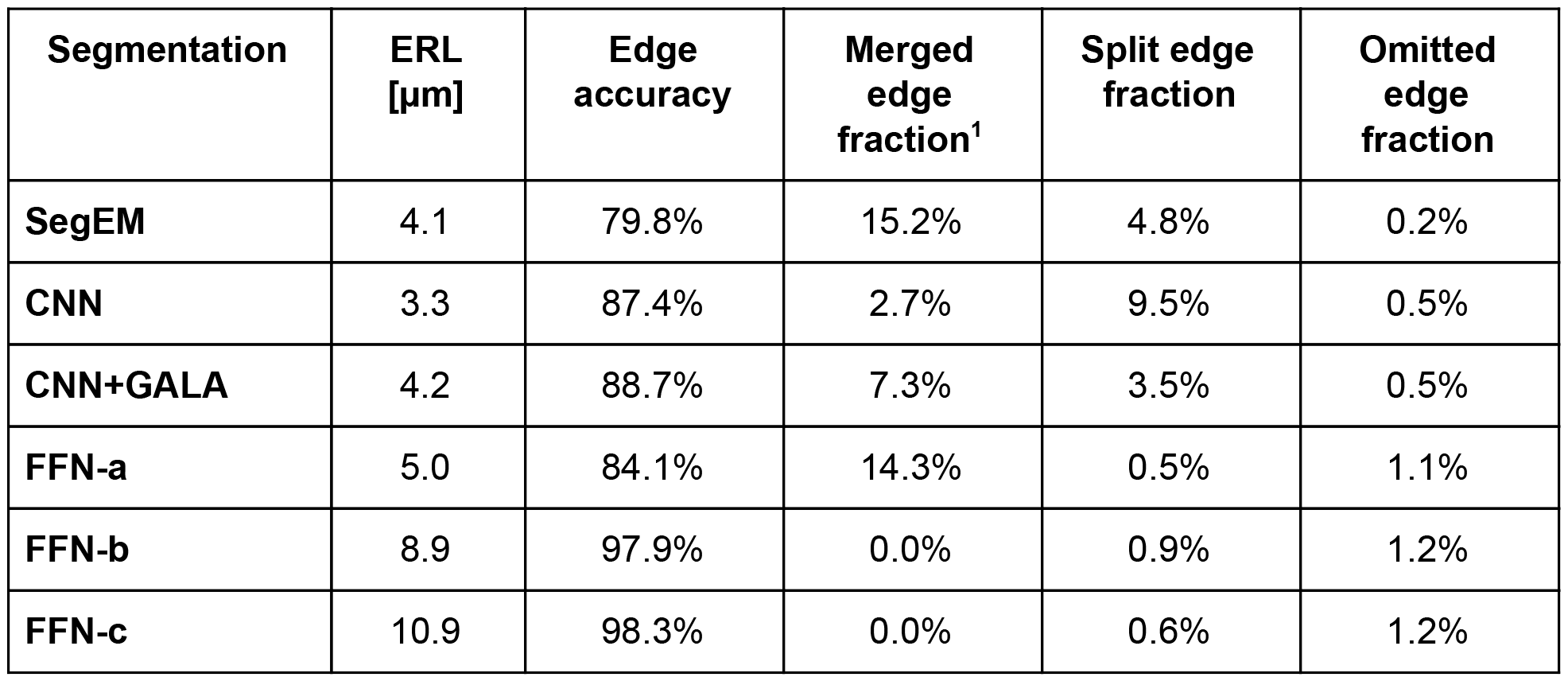
Evaluation of segmentation quality on the densely skeletonized [5 μm]^3^ subvolume.

**Supplementary Table 5.**
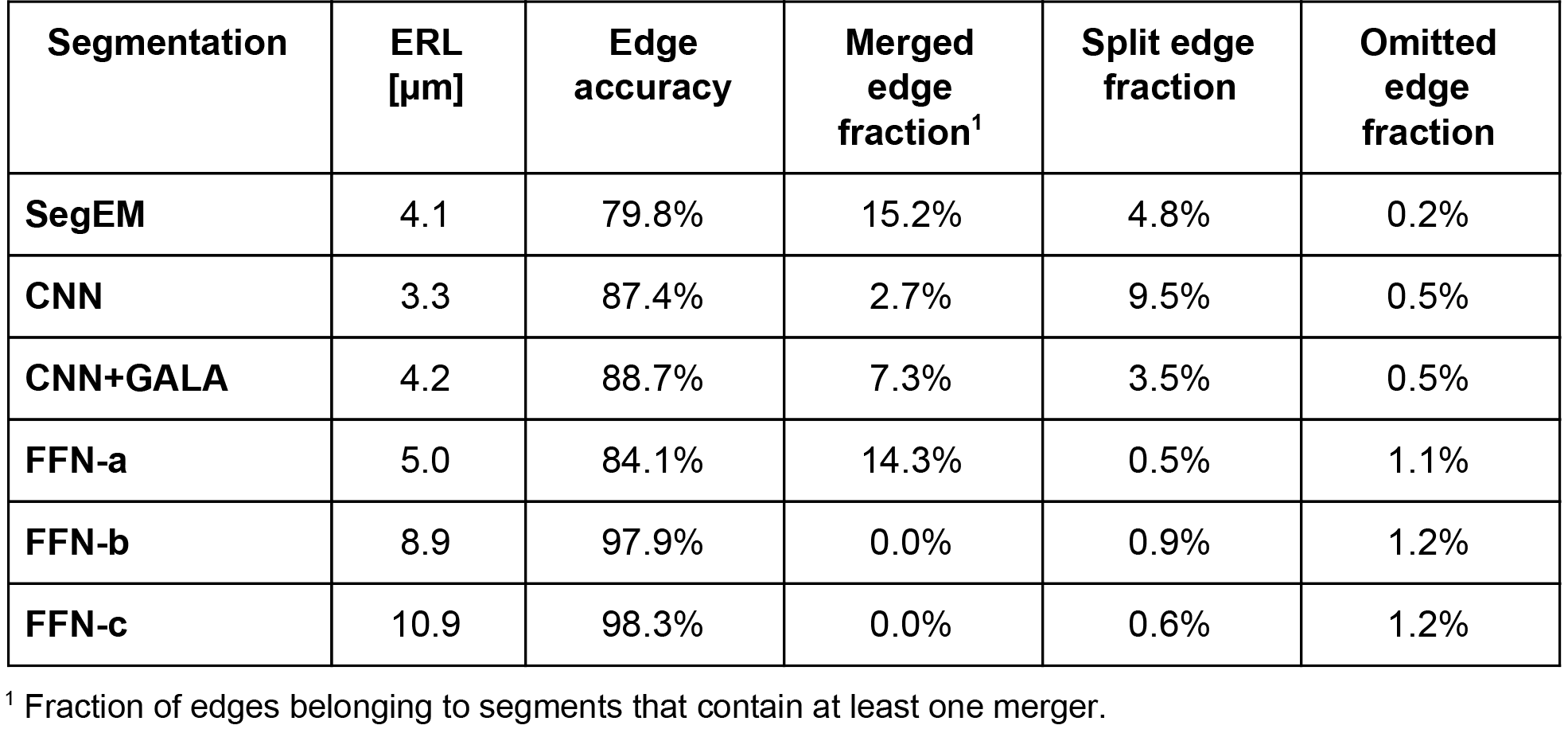
Evaluation of segmentation quality on the test set of 50 skeletons, with a total path length of 97 mm and max ERL of 2.1 mm.

Comparing the data in Sup. Tables 4 and 5, we observe that small-scale evaluations can severely underestimate the merged path length and the relative quality of different segmentations. For instance, in Sup. Tab. 4 FFN-c looks like a small incremental improvement over FFN-b, but when evaluated over the whole volume, the former is significantly better in terms of reconstructed error-free path length.

There are two main reasons for this. First, some data quality issues are inherently local to different parts of the volume, and hard to capture in a single small subvolume. For instance, the densely skeletonized subvolume does not suffer from any misaligned slices, cutting artifacts, or weakly stained neurites. Second, as the rate of errors in the segmentation decreases, getting good estimates of the different error rates requires sampling larger path lengths, at some point exceeding those contained within a small subvolume.

### Ablation Experiments

To elucidate the impact of the recurrent and single-object nature of the FFNs, we trained four different network variants:

1. boundary prediction with no recurrent channel (standard approach),
2. boundary prediction with recurrent channel (multi-object prediction),
3. memory-less FFN (single object prediction, no recurrent channel),
4. full FFN (recurrent single object prediction).

Only the minimum required changes were made between the experiments, and all other parameters, such as the architecture of the network were kept fixed.

For experiments 1) and 2) with boundary prediction networks, the training data was formed by first applying morphological erosion with radius 1 to the ground truth data to ensure separation of nearby neurites, and then binarizing the resulting image. The soft labels of 0.05 for background and 0.95 for object interior were used, similarly to the original FFN. After inference, the boundary network predictions were thresholded at 0.5, and converted into a segmentation by computing the connected components of the regions labeled as object interior.

For experiments 1) and 3), the second channel of the network input was set to a uniform empty image at value 0.05, and fixed-step movement procedure was used with a step size of (8, 8, 4) voxels. The “disconnected voxel bias” was active, allowing the network to override prior predictions of “object interior” to “exterior”, but not vice versa.

**Supplementary Table 6.**
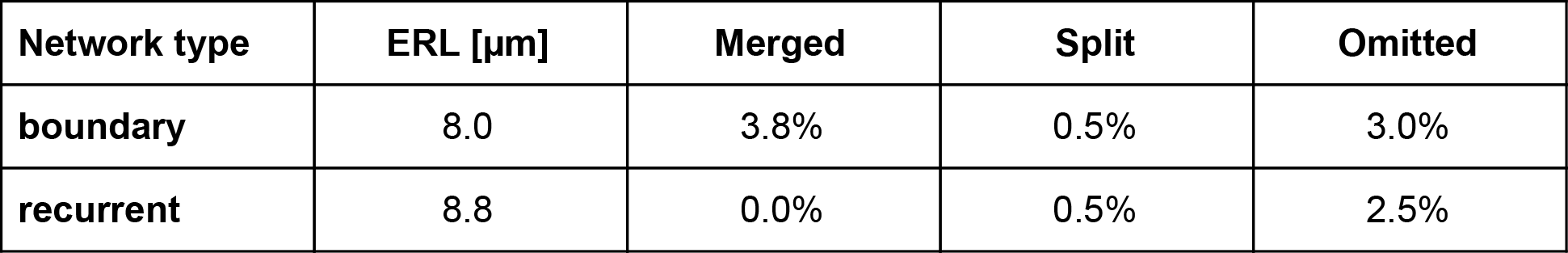
Evaluation of segmentation quality of different network types.

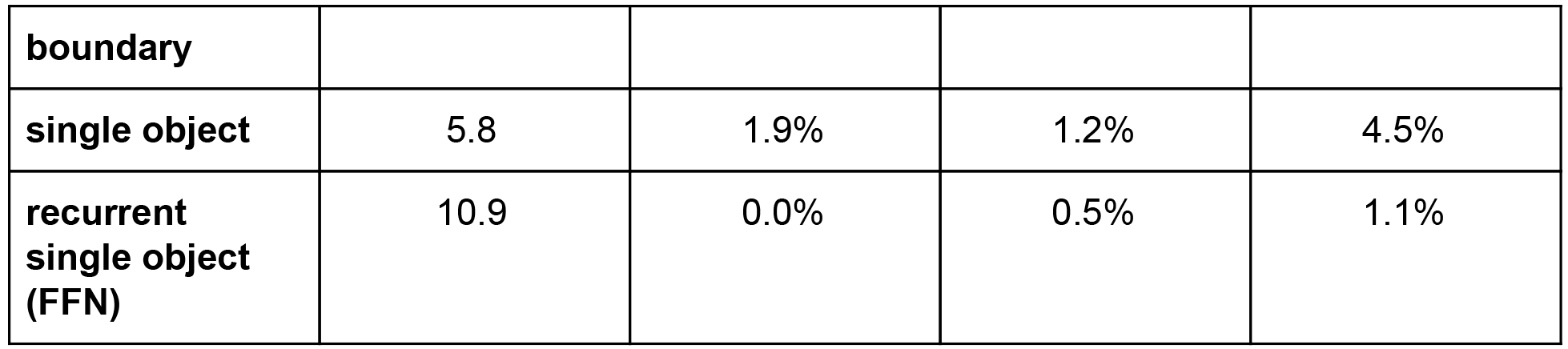

The results of our experiments presented in Sup. Table 6 suggest that the recurrent nature of the network is the main factor responsible for its segmentation accuracy, driven by its ability to eliminate mergers. These small scale experiments do not show the single object nature of the FFN to have a significant impact on the results. We note however, that this property is crucial for other procedures used in our pipeline (consensus, agglomeration), which were not applied here, but which were necessary to obtain good segmentations at the scale of the whole volume.

### Neuron Reconstructions in Skeleton Test Set

**Supplementary Figure 3.**
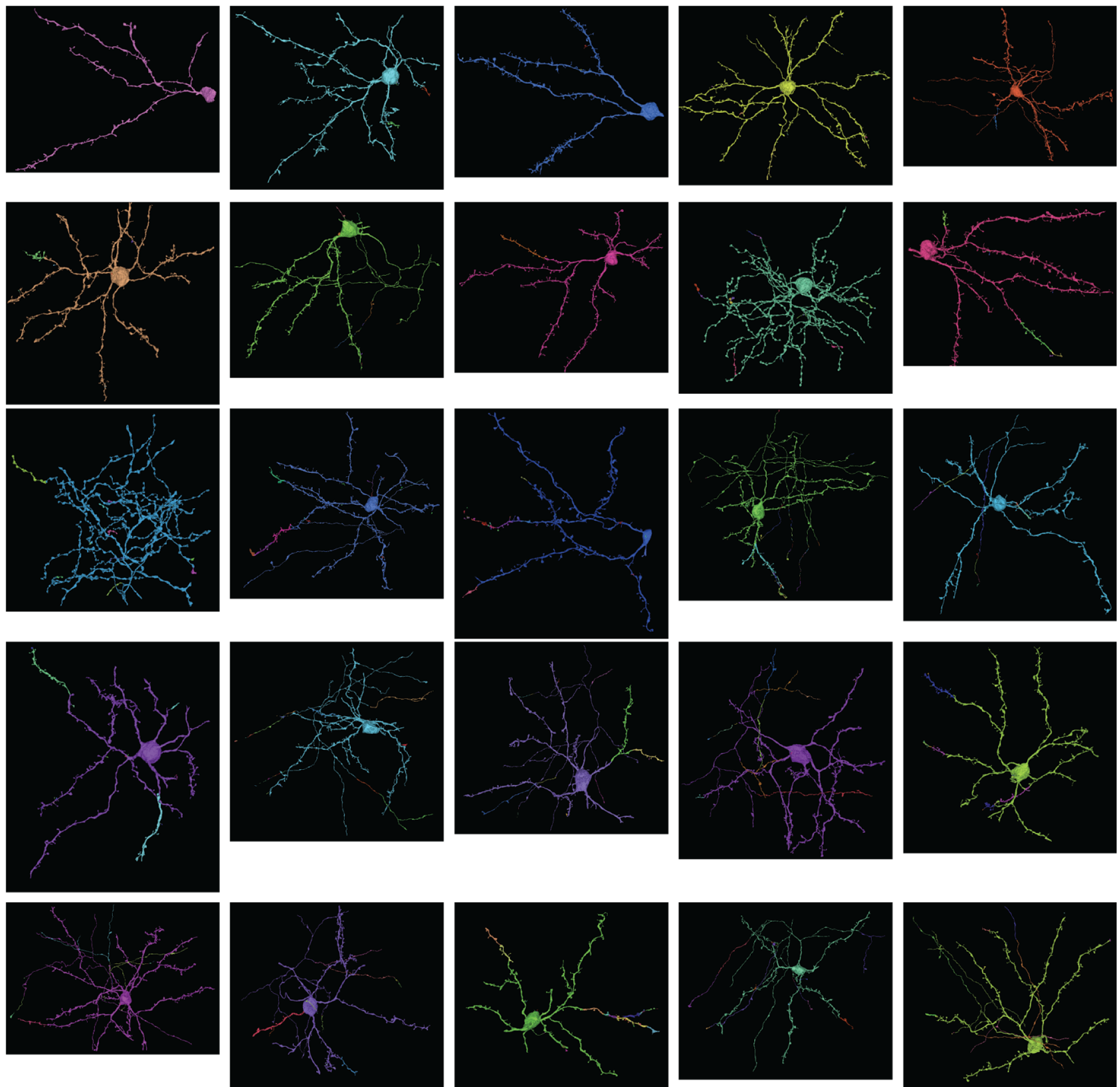
Qualitative analysis of segmentation accuracy. Different colors indicate different segments. Neurons reconstructed with the full pipeline (FFN-c) ordered from largest (92%, top-left) to shortest (0%, bottom-right) fraction of maximum expected run lengths according to the skeleton ground truth test set.

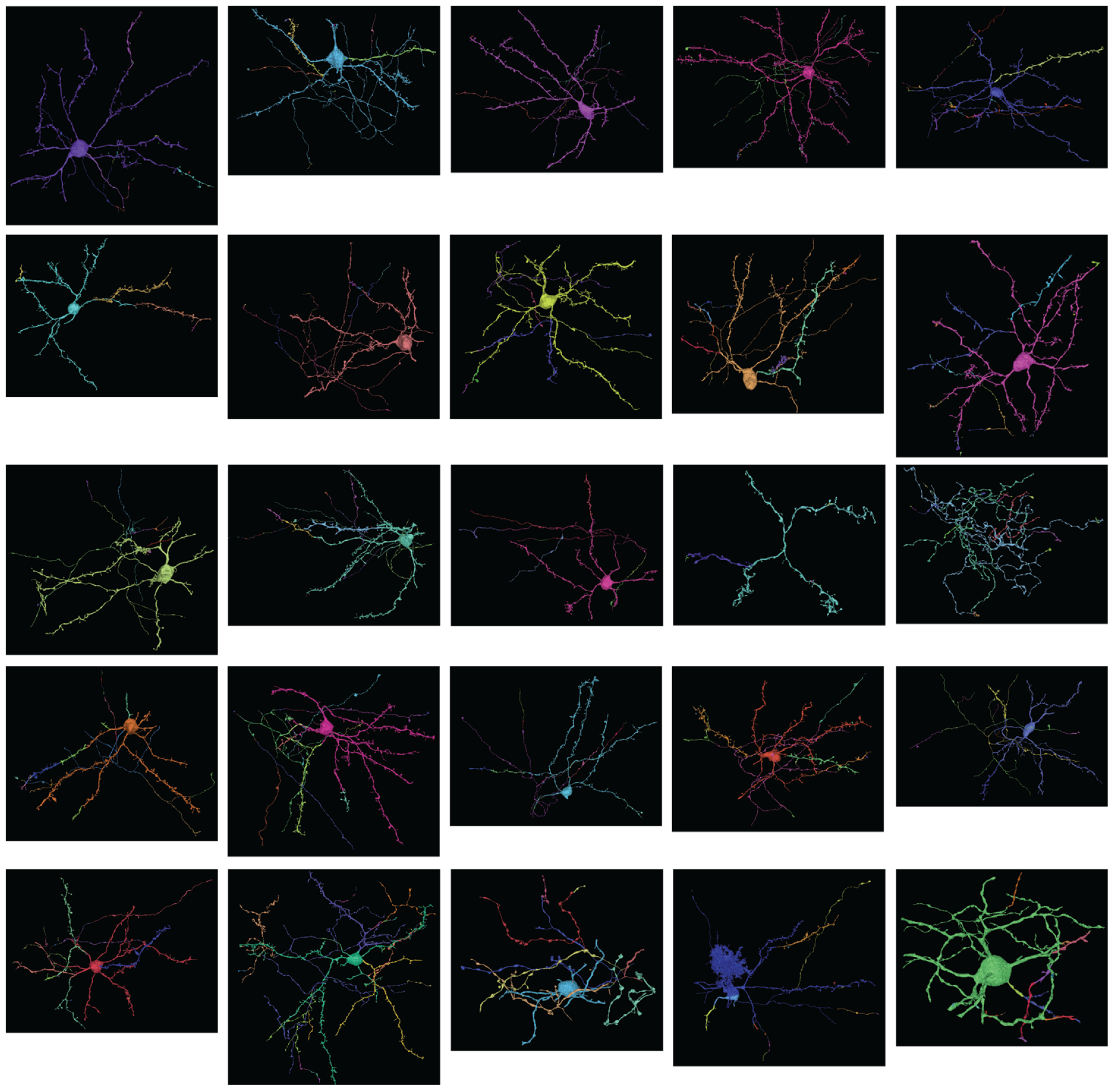

### FFN Reconstruction of Single Neurite

FILE ATTACHED: supplementary_video_1_ffn.mov

**Supplementary Video 1.** FFN reconstruction of a single neurite (i.e., seeded from a single voxel) in J0126 volume.

## References

1. Macagno, E. R., Levinthal, C. & Sobel, I. Three-dimensional computer reconstruction of neurons and neuronal assemblies. Annu. Rev. Biophys. Bioeng. 8, 323–351 (1979).

2. Harris, K. M., Jensen, F. E. & Tsao, B. Three-dimensional structure of dendritic spines and synapses in rat hippocampus (CA1) at postnatal day 15 and adult ages: implications for the maturation of synaptic physiology and long-term potentiation [published erratum appears in J Neurosci 1992 Aug; 12 (8): following table of contents]. Journal of Neuroscience 12, 2685–2705 (1992).

3. Ventura, R. & Harris, K. M. Three-dimensional relationships between hippocampal synapses and astrocytes. J. Neurosci. 19, 6897–6906 (1999).

4. Fiala, J. C. Reconstruct: a free editor for serial section microscopy. J. Microsc. 218, 52–61 (2005).

5. Briggman, K. L. & Bock, D. D. Volume electron microscopy for neuronal circuit reconstruction. Curr. Opin. Neurobiol. 22, 154–161 (2012).

6. Jain, V., Seung, H. S. & Turaga, S. C. Machines that learn to segment images: a crucial technology for connectomics. Curr. Opin. Neurobiol. (2010).

7. Kim, J. S. et al. Space-time wiring specificity supports direction selectivity in the retina. Nature 509, 331–336 (2014).

8. Helmstaedter, M., Briggman, K. L. & Denk, W. High-accuracy neurite reconstruction for high-throughput neuroanatomy. Nat. Neurosci. 14, 1081–1088 (2011).

9. Helmstaedter, M. et al. Connectomic reconstruction of the inner plexiform layer in the mouse retina. Nature 500, 168–174 (2013).

10. Cardona, A. TrakEM2: an ImageJ-based program for morphological data mining and 3d modeling. in Proceedings of the ImageJ User and Developer Conference (2006).

11. Berning, M., Boergens, K. M. & Helmstaedter, M. SegEM: efficient image analysis for high-resolution connectomics. Neuron 87, 1193–1206 (2015).

12. Plaza, S. M. Focused Proofreading to Reconstruct Neural Connectomes from EM Images at Scale. in Deep Learning and Data Labeling for Medical Applications 249–258 (Springer, Cham, 2016).

13. Jain, V. et al. Supervised Learning of Image Restoration with Convolutional Networks. in Computer Vision, 2007. ICCV 2007. IEEE 11th International Conference on 1–8 (2007).

14. Ciresan, D., Giusti, A., Schmidhuber, J. & Others. Deep neural networks segment neuronal membranes in electron microscopy images. in Advances in Neural Information Processing Systems 25 2852–2860 (2012).

15. Andres, B., Koethe, U., Helmstaedter, M., Denk, W. & Hamprecht, F. A. Segmentation of SBFSEM volume data of neural tissue by hierarchical classification. in DAGM 142–152 (Springer, 2008).

16. Kaynig, V. et al. Large-scale automatic reconstruction of neuronal processes from electron microscopy images. Med. Image Anal. 22, 77–88 (2015).

17. Knowles-Barley, S. et al. RhoanaNet Pipeline: Dense Automatic Neural Annotation. arXiv [q-bio.NC] (2016).

18. Beier, T. et al. Multicut brings automated neurite segmentation closer to human performance. Nat. Methods 14, 101–102 (2017).

19. Turaga, S. C., Briggman, K. L., Helmstaedter, M., Denk, W. & Seung, H. S. Maximin affinity learning of image segmentation. in NIPS (2009).

20. Jain, V. et al. Boundary Learning by Optimization with Topological Constraints. in Computer Vision and Pattern Recognition, IEEE Computer Society Conference on (2010).

21. Turaga, S. C. et al. Convolutional Networks Can Learn to Generate Affinity Graphs for Image Segmentation. Neural Comput. 22, 511–538 (2010).

22. Martin, D. R., Fowlkes, C. C. & Malik, J. Learning to detect natural image boundaries using local brightness, color, and texture cues. IEEE Trans. Pattern Anal. Mach. Intell. 530–549 (2004).

23. Funke, J., Andres, B., Hamprecht, F. A., Cardona, A. & Cook, M. Efficient automatic 3D-reconstruction of branching neurons from EM data. in 2012 IEEE Conference on Computer Vision and Pattern Recognition 1004–1011 (2012).

24. Goodfellow, I., Bengio, Y. & Courville, A. Deep Learning. (MIT Press, 2016).

25. Denk, W. & Horstmann, H. Serial block-face scanning electron microscopy to reconstruct three-dimensional tissue nanostructure. PLoS Biol. 2, e329 (2004).

26. Ronneberger, O., Fischer, P. & Brox, T. U-net: Convolutional networks for biomedical image segmentation. in International Conference on Medical Image Computing and Computer-Assisted Intervention 234–241 (Springer, 2015).

27. Nunez-Iglesias, J., Kennedy, R., Plaza, S. M., Chakraborty, A. & Katz, W. T. Graph-based active learning of agglomeration (GALA): a Python library to segment 2D and 3D neuroimages. Front. Neuroinform. 8, (2014).

28. Maitin-Shepard, J. et al. Combinatorial energy learning for image segmentation. arXiv preprint arXiv:1506.04304 (2015).

29. Saalfeld, S., Fetter, R., Cardona, A. & Tomancak, P. Elastic volume reconstruction from series of ultra-thin microscopy sections. Nat. Methods 9, 717–720 (2012).

30. Dorkenwald, S. et al. Automated synaptic connectivity inference for volume electron microscopy. Nat. Methods 14, 435–442 (2017).

31. Arganda-Carreras, I. et al. Crowdsourcing the creation of image segmentation algorithms for connectomics. Front. Neuroanat. 9, 142 (2015).

32. Pallotto, M., Watkins, P. V., Fubara, B., Singer, J. H. & Briggman, K. L. Extracellular space preservation aids the connectomic analysis of neural circuits. Elife 4, (2015).

33. Zlateski, A. & Seung, H. S. Image segmentation by size-dependent single linkage clustering of a watershed basin graph. arXiv preprint arXiv:1505.00249 (2015).

34. Han, S., Mao, H. & Dally, W. J. Deep Compression: Compressing Deep Neural Networks with Pruning, Trained Quantization and Huffman Coding. arXiv [cs.CV] (2015).

35. Abadi, M. et al. TensorFlow: Large-Scale Machine Learning on Heterogeneous Distributed Systems.

36. LeCun, Y. et al. Backpropagation Applied to Handwritten Zip Code Recognition. Neural Comput. 1, 541–551 (1989).

37. LeCun, Y. A., Bottou, L., Orr, G. B. & Müller, K.-R. Efficient BackProp. in Neural Networks: Tricks of the Trade 9–48 (Springer, Berlin, Heidelberg, 2012).

38. Tschopp, F. Efficient Convolutional Neural Networks for Pixelwise Classification on Heterogeneous Hardware Systems. arXiv [cs.CV] (2015).

39. He, K., Zhang, X., Ren, S. & Sun, J. Identity mappings in deep residual networks. arXiv preprint arXiv:1603.05027 (2016).

40. He, K., Zhang, X., Ren, S. & Sun, J. Deep residual learning for image recognition. arXiv preprint arXiv:1512.03385 (2015).

